# mtDNA eQTLs and the m^1^A 16S rRNA modification explain mtDNA tissue-specific gene expression pattern in humans

**DOI:** 10.1101/495838

**Authors:** T. Cohen, C. Mordechai, A. Eran, D Mishmar

## Abstract

Expression quantitative trait loci (eQTLs) are instrumental in genome-wide identification of regulatory elements, yet were overlooked in the mitochondrial DNA (mtDNA). By analyzing 5079 RNA-seq samples from 23 tissues we identified association of ancient mtDNA SNPs (haplogroups T2, L2, J2 and V) and recurrent SNPs (mtDNA positions 263, 750, 1438 and 10398) with tissue-dependent mtDNA gene-expression. Since the recurrent SNPs independently occurred in different mtDNA genetic backgrounds, they constitute the best candidates to be causal eQTLs. Secondly, the discovery of mtDNA eQTLs in both coding and non-coding mtDNA regions, propose the identification of novel mtDNA regulatory elements. Third, we identified association between low m^1^A 947 MT-RNR2 (16S) rRNA modification levels and altered mtDNA gene-expression in twelve tissues. Such association disappeared in skin which was exposed to sun, as compared to sun-unexposed skin from the same individuals, thus supporting the impact of UV on mtDNA gene expression. Taken together, our findings reveal that both mtDNA SNPs and mt-rRNA modification affect mtDNA gene expression in a tissue-dependent manner.

## Introduction

The mitochondrion is the major source of cellular energy (ATP) and a pivotal player in cell life and death, and hence is critical for the life of most tissues. However, mitochondrial mass, morphology and the amount of ATP produced via the oxidative phosphorylation system (OXPHOS) differ between tissues (Fernandez-Vizarra et al. 2011). Accordingly, tissue-dependent mitochondrial disorders are frequently reported (Boczonadi et al. 2018), and mitochondrial DNA (mtDNA) common variants often associate with altered tendency to develop tissue-dependent phenotypes (Marom et al. 2017). As mitochondrial activity differs among tissues, it is reasonable that regulation of such functions will be tissue-dependent.

Although most mitochondrial factors are encoded by the nuclear genome (nDNA) (Calvo et al. 2016), 37 essential genes are encoded by the mtDNA in all studied vertebrates (including humans). Specifically, the human mtDNA encodes 13 protein subunits of the OXPHOS system, 22 tRNAs and two rRNAs (MT-RNR1 and MT-RNR2), which co-transcribed in 2-3 strand-specific polycistrons(Aloni and Attardi 1971). Recently we demonstrated co-regulation of mtDNA and nDNA-encoded OXPHOS genes across many human body sites (Barshad et al. 2018a). This analysis enabled identifying candidate regulatory factors, which both preferentially bind upstream regulatory elements of nDNA-encoded OXPHOS genes, and bind the human mtDNA *in vivo* (Blumberg et al. 2014). As accumulating evidence suggest that quite a few known regulators of nDNA genes’ transcription are imported into the mitochondria, bind specific regions throughout the mtDNA and regulate mtDNA transcription (Barshad et al. 2018b), we hypothesized that there are additional (and yet unknown) regulatory elements of mtDNA gene expression, which are not necessarily confined to the non-coding mtDNA control region.

Expression quantitative trait loci (eQTLs) have been widely used in genome-wide screens for regulatory elements (Westra and Franke 2014), yet were overlooked in the mtDNA. Previously, we and others performed in vitro transcription (Suissa et al. 2009) and RNA-seq in cell culture from multiple individuals to assess the impact of human mtDNA variants on mtDNA transcription (Cohen et al. 2016). Such experiments, however, assessed the impact of mtDNA variants on gene expression only in lymphoblastoid cells. Although an assessment of mtDNA gene expression in cytoplasmic cell lines (cybrids) enabled comparison of the impact of different mtDNA genetic backgrounds (haplogroups) on gene expression while controlling for the cell nucleus (Gomez-Duran et al. 2010; Kenney et al. 2014), these experiments were again performed using specific cell types, and specific mtDNA genetic backgrounds. As such, these efforts failed to explain how, and if, mtDNA gene expression differs among tissues (Fernandez-Vizarra et al. 2011). With this in mind, we recently analyzed RNA-seq experiments from 48 different human tissues, as well as single-cell data from different cell types in human and mouse brains, and found tissue and cell type variability in terms of mitochondrial-nuclear genes’ co-expression (Barshad et al. 2018a). Therefore, although tissue variability has been shown in terms of nDNA and mtDNA gene expression levels (Sudmant et al. 2015), it remains unclear whether the functional impact of inherited mitochondrial variants is influenced by tissue and/or cell types.

Here, we tested the association between mtDNA genetic variants and gene expression patterns in a variety of tissues, thereby identifying mitochondrial expression quantitative loci (mt-eQTLs). Specifically, we analyzed the mtDNA-encoded transcript sequences of samples collected by the GTEx consortium and assessed the association of mtDNA-encoded RNA sequence variants with altered gene expression. These analyses revealed that both ancient mtDNA haplogroup-defining variants and recurrent variants may act as eQTLs in a tissue-dependent manner. We also found that the low level of RNA-DNA difference at mtDNA position 2617, indicative of m^1^A 947 MT-RNR2 (16S) rRNA modification, was associated with differential mtDNA gene expression in several tissues, thus connecting mtDNA transcription and translation. The implications of these findings for the identification of novel putative mtDNA regulatory elements are discussed.

## Results

### Identification of tissue-dependent mtDNA eQTLs

To identify mt-eQTLs, we analyzed 5079 GTEx RNA-seq samples from 23 tissues, each surveyed in at least 100 individuals (Fig. 1). Since the entire human mtDNA sequence is transcribed, we used the RNA-seq data to reconstruct the personal mitochondrial genome of each sample, call variants, and assign each individual to an mtDNA-determined genetic background (i.e., haplogroups) (Kloss-Brandstatter et al. 2011) (Supplemental Table S1). To avoid sample size issues, we excluded samples with Asian mtDNA haplogroups from further analyses (haplogroups A, B, C, D and N1a9), due to their overall small representation (N=11). Notably, although the sequencing read coverage across the mtDNA differed among the tested body sites (Supplemental Fig. S1, it did not preferentially impact any certain mtDNA genetic backgrounds. This left us with sample sizes ranging from N=97 (ovary) to N=441 (whole blood) (Supplemental Table S1). We next assessed the association of each recorded mtDNA SNP with the expression level of mtDNA-encoded transcripts in tissues with a sample size of at least 250 individuals (N=8 tissues; Supplemental Table S2). Notably, to avoid erroneous signals coming from mtDNA fragments that were transferred to the cell nucleus during evolution (i.e., nuclear mitochondrial pseudogenes - NUMTs) (Mishmar et al. 2004), we employed unique mapping. This analysis revealed that seven ancient, haplogroup-defining SNPs (Levin et al. 2013) associated with altered mtDNA-encoded gene expression patterns in six of the tested tissues (Fig. 2, Table 1, Supplemental Dataset S1). Some of the identified SNPs associated with increased expression of all tested mtDNA-encoded genes (e.g., T2 haplogroup in skeletal muscle, V haplogroup in tibial artery) or with the altered expression of a number of mtDNA-encoded genes (Table 1). Nevertheless, all identified mtDNA SNP associations with mtDNA gene expression were specific to particular body sites, such as the association of haplogroup L2-defining SNPs with increased expression of specific mtDNA genes in the ovary and tibial nerve.

**Fig. 1:**
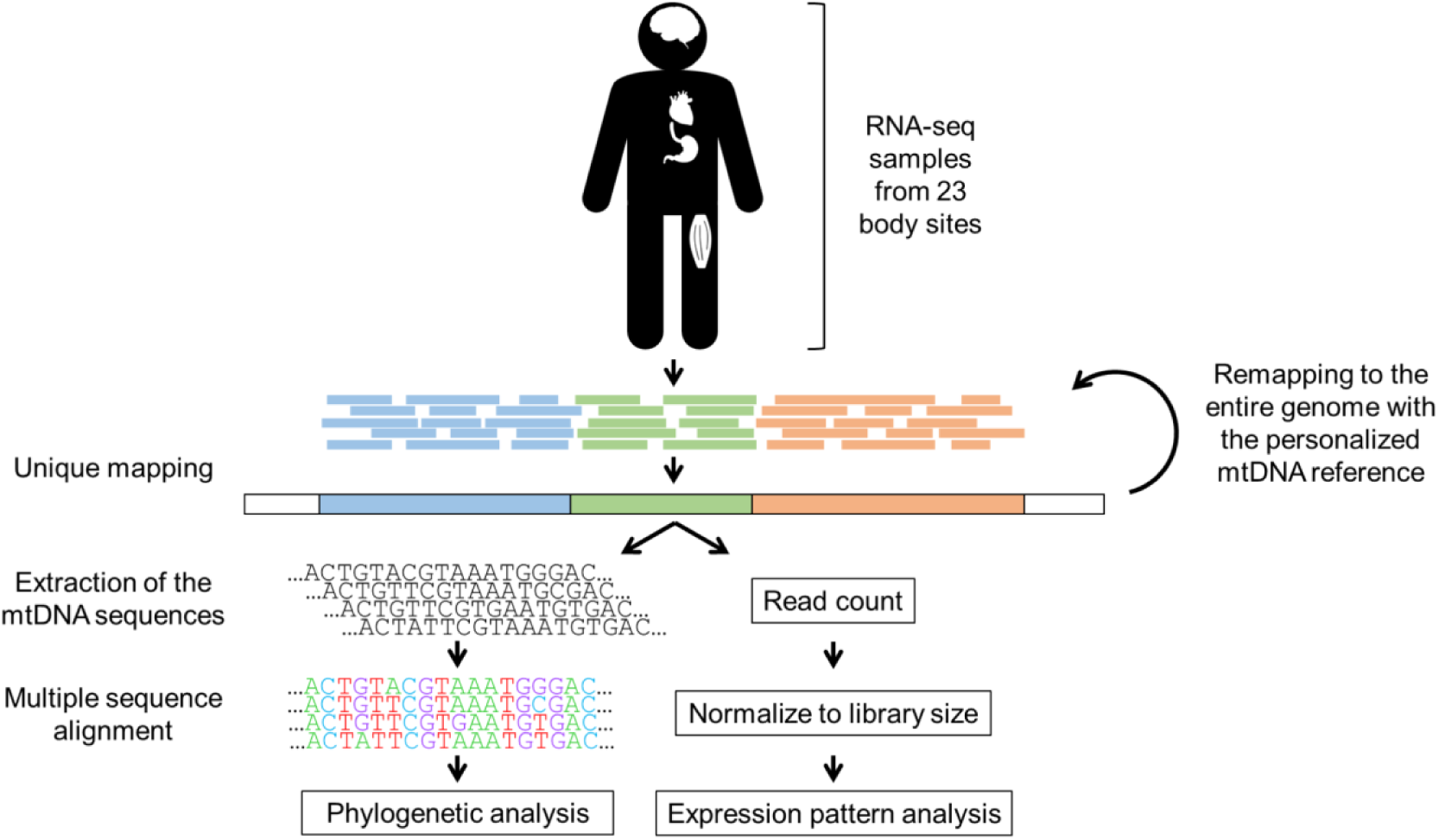
Study design. RNA-seq reads from 23 body sites were mapped against the human genome (GRCh38), allowing output of reads that mapped to single loci (unique mapping). The extracted mtDNA sequences were used to perform phylogenetic analysis and as specific reference sequences for each sample in a remapping process. After remapping, a second phylogenetic analysis was performed, and reads were counted and normalized to library size, allowing for testing of the expression pattern of each sample according to its phylogenetic data.

**Fig. 2:**
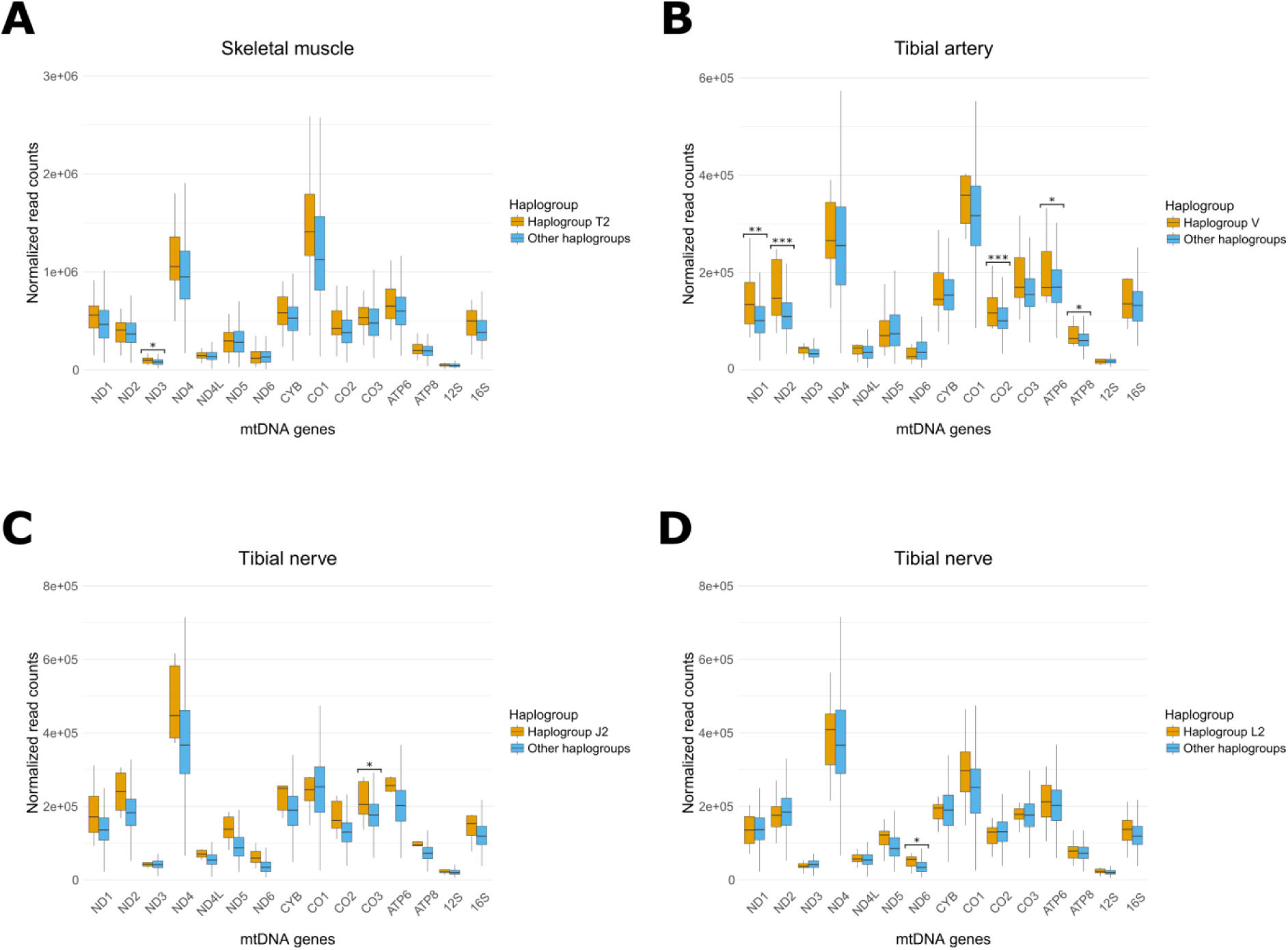
A screen of all identified mtDNA SNPs highlights association of haplogroup-defining SNPs with mtDNA gene expression at specific body sites. An unbiased mtDNA SNP by SNP association study revealed that expression levels of mtDNA genes were higher (orange) in individuals belonging to haplogroups T2 (A), V (B), J2 (C) and L2 (D), when compared to the rest of the cohort (blue). X axis-mtDNA genes. Y axis-normalized read counts. (*) < 1.19 × 10^−5^, (**) < 1 × 10^−6^, (***) < 1 × 10^−7^.

**Table 1:** mtDNA SNPs associate with altered mtDNA gene expression. Arrow indicates elevated (upward) or reduced (downward) mtDNA gene expression levels.

| SNP (Haplogroup) | Tissues | Significantly affected genes |
| --- | --- | --- |
| C7476T, G15257T (J2) | Tibial nerve (6/303) | MT-CO3 ↑ |
| G8206A (L2) | Tibial nerve (17/303)<br>Unexposed skin (11/245)<br>Ovary (5/96) | MT-ND6 ↑<br>MT-CO3, MT-ATP8 ↑<br>MT-ND5, ND6 ↑ |
| A11812G, A14233G (T2) | Skeletal muscle (28/410)<br>Cerebellum (6/129)<br>Cerebral hemisphere (5/107) | MT-ND3 ↑<br>All mtDNA genes ↓<br>All mtDNA genes ↑ |
| G4580A (V) | Tibial artery (11/317)<br>Cortex (6/118) | MT-ND1, MT-ND2, MT-CO2, MT-ATP6, MT-ATP8 ↑<br>All mtDNA genes ↑ |
| A7424G (L3d) | Adipose (5/321) | All MT-ND genes, MT-CYB, MT-CO1, MT-RNR2 ↑ MT-CO3 ↓ |
| G7337A (L1c2) | Skeletal muscle (6/410)<br>Sun-exposed skin (6/348) | All mtDNA genes ↑<br>MT-RNR1 ↑ |

To assess mtDNA SNP association with mtDNA gene expression at body sites represented by smaller sample sizes, we next considered the list of SNPs that associated with mtDNA-encoded gene expression in any of the above-mentioned eight tissues with larger sample sizes (Tables 1 and 2). Such analysis revealed five additional body sites (i.e., sun-unexposed skin, cerebellum, cortex, cerebral hemisphere and ovary) in which mtDNA gene expression associated with SNPs defining the J2, L2, T2 and V haplogroups (Fig. 3, Table 1). Interestingly, although distinct mtDNA gene expression pattern was associated with the T2 haplogroup in both the cerebellum and the cerebellar hemisphere, the mtDNA transcripts were decreased in the former, but increased in the latter, probably due to the differential neuronal (and non-neuronal) cell composition in these two types of samples (Table 1). Notably, samples from different tissues originating from the same individuals presented different mtDNA SNP associations with mtDNA gene expression (Supplemental Dataset S1). As the mtDNA is considered a single locus in full linkage disequilibrium, and hence, all mtDNA SNPs in a given individual are in full linkage, our results support tissue-dependent association of mtDNA genetic backgrounds with mtDNA gene expression.

**Table 2:**
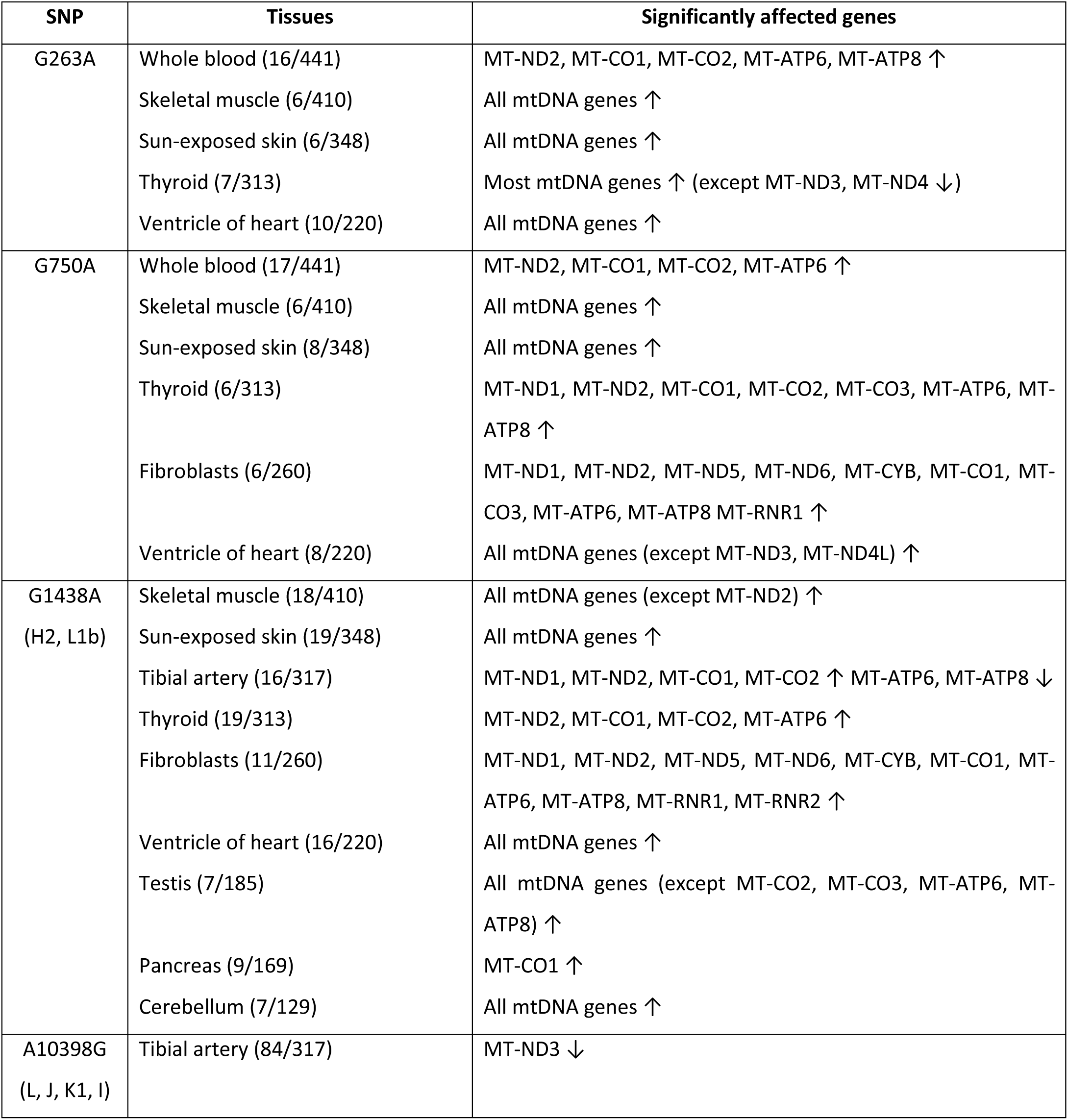
Recurrent mt-eQTLs. Arrow indicates elevated (upward) or reduced (downward) mtDNA gene expression levels.

**Fig. 3:**
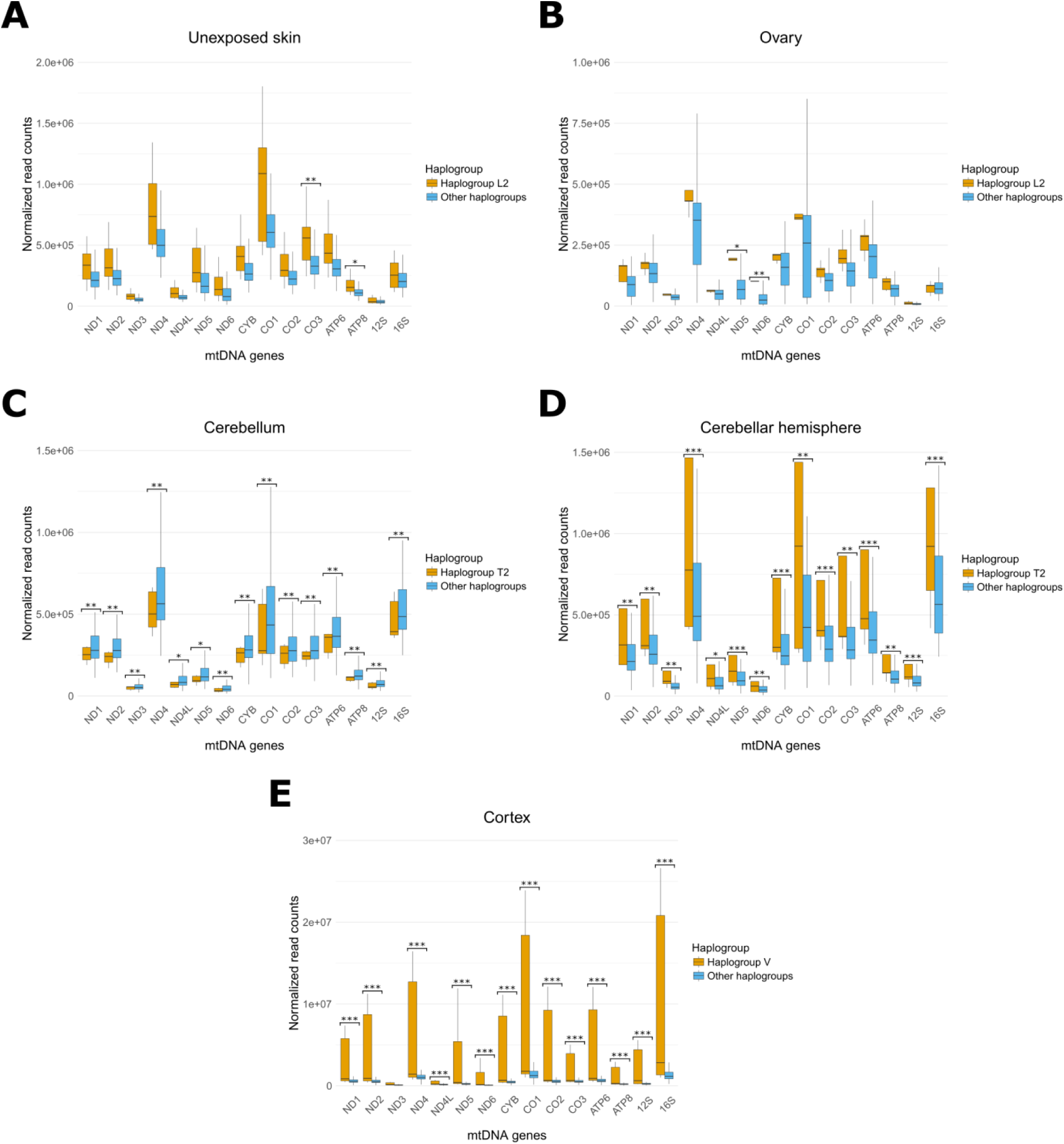
Analysis of selected mtDNA SNPs unravels their association with mtDNA gene expression at additional body sites. Expression levels of mtDNA genes were higher (orange) in individuals belonging to haplogroup L2 in the skin which was not exposed to sun (A), and in the ovary (B). In individuals belonging to haplogroups T2 (orange) mtDNA genes expression was lower in the cerebellum (C) but higher in the cerebellar hemisphere (D). Finally, in individuals belonging to haplogroup V (orange) mtDNA gene expression was higher in the cortex (E). Notably, all comparisons were performed against individuals from the rest of the cohort (blue). X axis-mtDNA genes. Y axis-normalized read counts. (*) < 4.17 × 10^−4^, (**) < 1 × 10^−4^, (***) < 1 × 10^−5^.

### Identification of recurrent mt-eQTLs

The SNPs we identified thus far as mtDNA eQTLs are part of haplogroups. As such, one cannot distinguish between their putative direct (i.e. causal) or indirect association (i.e. linkage with other variants) with mtDNA gene expression. However, in addition to the observed association of mtDNA gene expression with ancient haplogroup-defining SNPs, we also identified association of altered mtDNA gene expression with four mtDNA SNPs that reoccurred during the course of human evolution (Levin et al. 2013). Specifically, the transition at mtDNA position 750, which occurs in multiple haplotypes, associated with differential expression in 6 tissues (Table 2). Similarly, the transition at mtDNA position 263, which is shared by individuals assigned to a variety of mitochondrial haplogroups, associated with mtDNA gene expression in 5 of the 6 tissues that showed association with the 750 SNP (Table 2). A transition in mtDNA position 1438, which re-occurred in the mtDNA branches leading to the H2 and L1b haplogroups (Levin et al. 2013), associated with altered mtDNA gene expression at 9 body sites (Table 2). Finally, the transition at mtDNA position 10398, which reoccurred in the L, J, I and K1 haplogroups (Levin et al. 2013), associated with differential expression of the ND3 gene in the tibial artery. As recurrent mtDNA SNPs have previously been associated with phenotypes independent of their associated haplotypes (Levin et al. 2013; Levin and Mishmar 2017), such markers can be considered as causal eQTLs, which likely reside within or adjacent to regulatory elements of mtDNA gene expression.

### nDNA SNPs and co-expression analyses offer candidates that partially explain the association of mtDNA SNPs with mtDNA gene expression

All known regulatory factors of mtDNA gene expression are encoded by the nDNA (Barshad et al. 2018b). Thus, we asked whether SNPs within 116 genes that participate in mitochondrial transcriptional and post-transcriptional regulation (Cohen et al. 2016) (Supplemental Table S3) could explain the identified mtDNA eQTLs. Although we detected association of certain nDNA SNPs with mtDNA gene expression (Supplemental Dataset S2), apart from a weak association of a missense mutation in MRPS17 (i.e. Mitochondrial Ribosomal Protein S17) in the context of the SNP at mtDNA position 1438, none of the analyzed nDNA SNPs explained mtDNA haplogroup association with gene expression, likely due to the small sample sizes of each of the analyzed haplogroups.

Additionally, we assessed co-expression of the above-mentioned 116 nDNA genes with mtDNA gene expression in the context of our identified mt-eQTLs. Firstly, while considering haplogroup-associated eQTLs, the only significant co-expression was of ALKBH7 (i.e., mitochondrial Alpha-Ketoglutarate-Dependent Dioxygenase AlkB Homolog 7) with all mtDNA-encoded genes in skin which was not exposed to sun, in individuals belonging to the L2 haplogroup (Supplemental Dataset S3). Nevertheless, such association was not identified in other tissues that highlighted association of L2 haplogroups with mtDNA gene expression. This suggests that the potential contribution of ALKBH7 to the L2 haplogroup associated mt-eQTL is minor.

Secondly, while considering the recurrent SNPs at positions 750 and 1438 none of the identified nDNA co-expressed genes explained these mt-eQTLs in more than two body sites (Supplemental Dataset S3, Supplemental Table S4). Since these eQTLs associated with mtDNA gene expression in 6 and 9 body sites, respectively, the nDNA genes could only partially explain such association. In contrast, we found that the expression of C21orf33 (i.e., Glutamine Amidotransferase Like Class 1 Domain Containing 3A) associated with the mt-eQTL at position 263 in 4 out of 5 body sites, which displayed the impact of position 263 on mtDNA gene expression. Similarly, we found co-expression of AK4 (i.e., mitochondrial Adenylate Kinase 4) with the mt-eQTL at mtDNA position 10398 in the tibial artery (e.g., the only body site in which mtDNA gene expression associated with this mtDNA position, Supplemental Dataset S3, Supplemental Table S4). These pieces of evidence may offer candidate regulators of mtDNA gene expression in the context of mtDNA recurrent SNPs.

### An m^1^A 947 16S rRNA modification associates with altered mtDNA gene expression at multiple body sites

RNA-seq reads enable the identification of RNA-specific sequence changes (i.e., RNA-DNA differences), which in many cases represent signatures of specific RNA modifications (Bar-Yaacov et al. 2013; Bar-Yaacov et al. 2016). Accordingly, we asked whether mitochondrial RNA-specific changes also associate with altered mtDNA gene expression. Previously, we found that mtDNA position 2617 (Bar-Yaacov et al. 2013), reported to be invariant in the mtDNA sequences of nearly 10,000 individuals representing all major global populations (Bar-Yaacov et al. 2013), instead exhibited a mixture of adenine, thymidine and occasionally guanine-containing reads in RNA-seq of all tissues studied thus far (Bar-Yaacov et al. 2013) and in many individuals (Hodgkinson et al. 2014). Recently, we found that such base-mixture echoes an m^1^A modification of 16S rRNA transcript position 947 (m^1^A 947) (Bar-Yaacov et al. 2013; Bar-Yaacov et al. 2016). Firstly, there was considerable variation in the levels of the m^1^A 947 16S rRNA modification among different body sites (Fig. 4). Specifically, the modification levels ranged from the lowest in whole blood to the highest in both the atrial appendage (heart) and the thyroid. Second, a bimodal distribution of the samples emerged, dividing the tested individuals into those with a lower modification level (i.e., <45% thymidine-harboring reads at mtDNA position 2617) versus those with a higher level of m^1^A 947 (i.e., >55% thymidine-harboring reads at the same position). As such, the level of m^1^A 947 16S rRNA modification cannot be explained as being a strictly continuous trait (Fig. 4). We then found that this modification associates with mtDNA gene expression at multiple body sites (Fig. 5, Supplemental Fig. S2). Notably, most body sites that showed mtDNA gene expression association with the m^1^A 947 16S rRNA modification (i.e., tibial artery, thyroid, skin not exposed to sun, atrial appendage of the heart, mammary tissue, stomach, colon, pancreas, frontal brain cortex and the ovary) presented lower gene expression in samples with higher modification levels (i.e. >55% of transcripts) (Fig. 5, and Supplemental Fig. S2). At the same time, the caudate and whole blood displayed an opposite effect (Fig. 5A and Supplemental Fig. S2). As m^1^A 947 16S rRNA modification likely affects mitochondrial protein translation (Bar-Yaacov et al. 2016), our results suggest an impact of modified mito-ribosome levels on mtDNA gene expression, consistent with cross-talk between mtDNA transcription and translation.

**Fig. 4:**
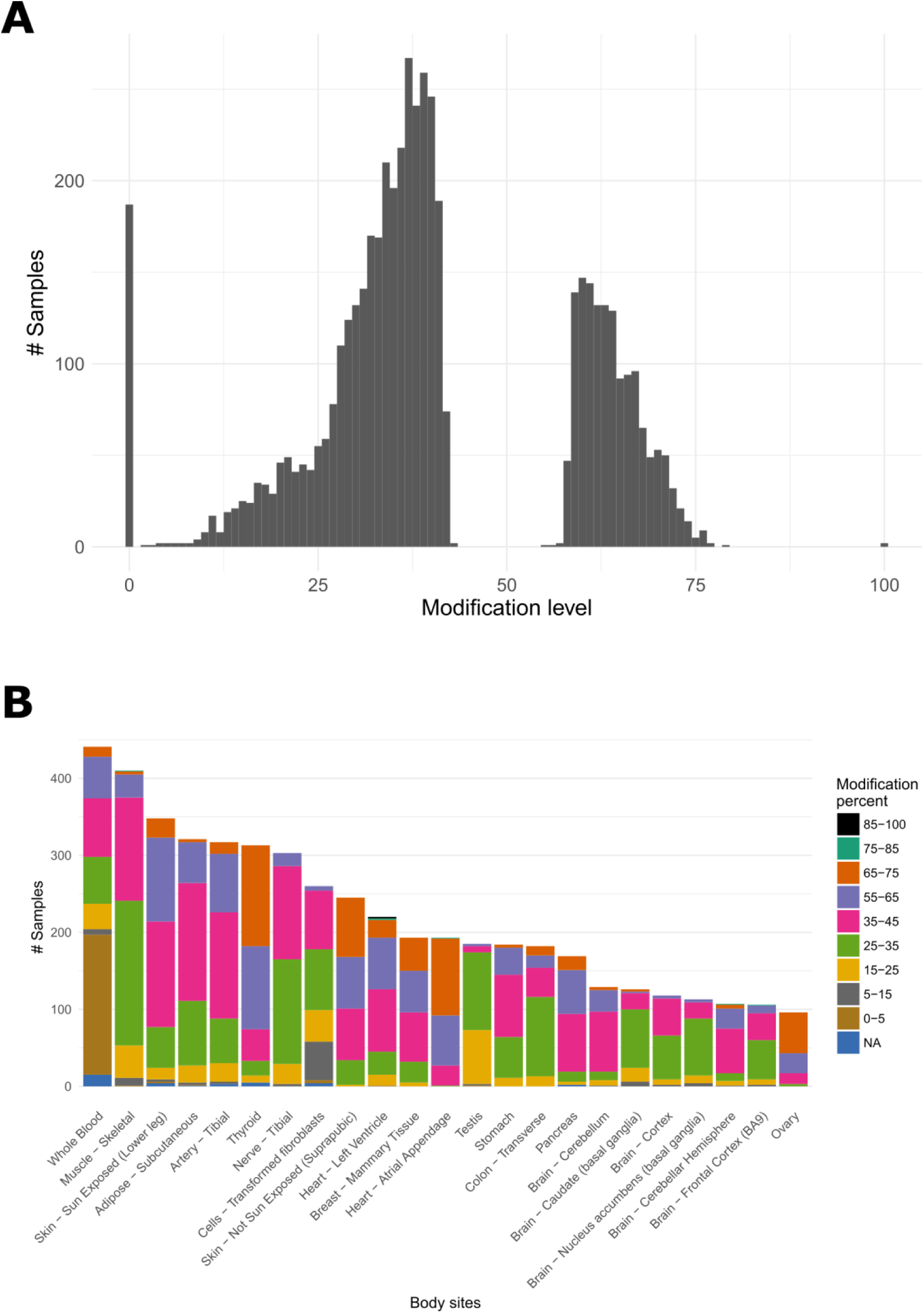
The distribution of m^1^A 947 16S rRNA modification levels across body sites and samples. (A) m^1^A 947 16S rRNA modification levels among samples showed bi-modal distribution. X axis-modification level in percentages. Y axis-number of samples. (B) Distribution of m^1^A 947 16S rRNA modification levels across body sites. X axis-body sites. Y axis-number of samples presenting a given modification percentage.

**Fig. 5:**
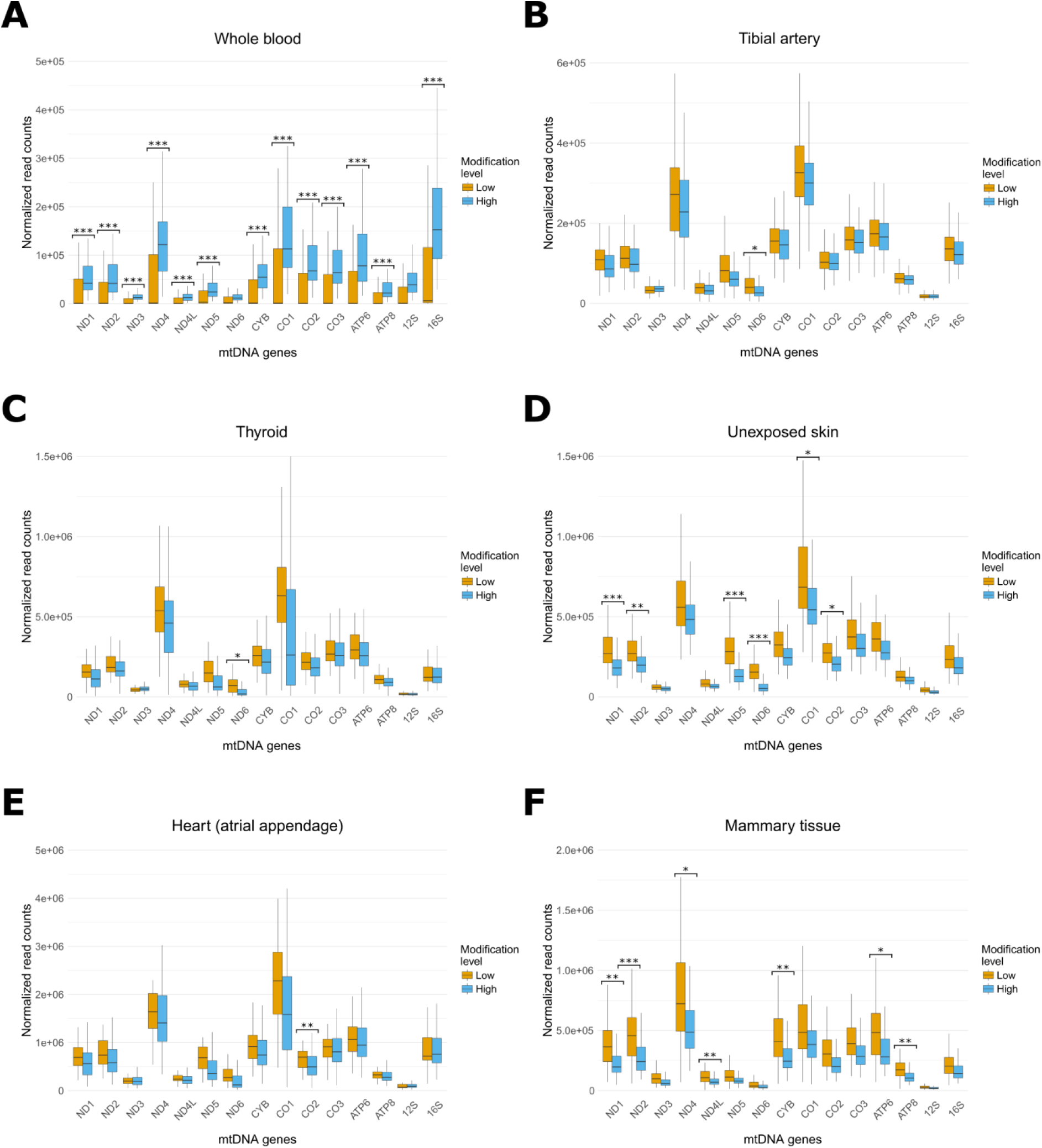
mtDNA gene expression patterns differ between individuals with low vs. high m^1^A 947 16S rRNA modification levels. Shown are representative body sites which display differential mtDNA gene expression in association with the m^1^A 947 16S rRNA modification levels. The results in the full set of analyzed body sites are displayed in Supplemental Fig. S2. Orange-samples with a low modification level; Blue - samples with a high modification level. mtDNA gene expression levels were higher in individuals with high modification levels in whole blood (A), and lower in the tibial artery (B), thyroid (C), sun-unexposed skin (D), atrial appendage of the heart (E) and in the mammary tissue (F). X axes-mtDNA genes. Y axes-normalized read counts. Panels a-c: (*) < 1.19 × 10^−5^, (**) < 1 × 10^−6^, (***) < 1 × 10^−7^. Panels d-f: (*) < 4.17 × 10^−4^, (**) < 1 × 10^−4^, (***) < 1 × 10^−5^.

### TRMT61B only partially explains the association of m^1^A 947 16S rRNA modification with altered mtDNA gene expression

We next sought to define the best candidate nDNA factors responsible for modulating the association of the m^1^A 947 16S rRNA modification with mtDNA gene expression. To this end, we screened the results from RNA-seq experiments involving the twelve tissues in which such modification associated with mtDNA gene expression for co-expressing nDNA-encoded factors, taking into account where high versus low levels of modification were observed. Firstly, we performed co-expression analysis to screen for candidate regulators of the discovered association of mtDNA gene expression with the m^1^A 947 16S rRNA modification levels in the above-mentioned 12 body sites (Supplemental Table S5). Although 107 out of the 116 candidate mtDNA gene expression regulators displayed significant co-expression values (p<4.3 × 10^−4^, Pearson correlation after Bonferroni corrections), none of these candidates had a significant co-expression value in more than 7 of the 12 tested body sites. Notably, we noticed that SLIRP was the only gene candidate that co-expressed, and displayed slightly higher expression level along with the modified 16S rRNA transcripts (i.e., higher levels of T-containing reads) in whole blood and the caudate (Supplemental Table S5). Secondly, TRMT61B, an RNA methyltransferase recently shown to introduce the m^1^A 947 16S rRNA modification (Bar-Yaacov et al. 2016), significantly co-expressed with mtDNA genes only in the whole blood and in skin that was not exposed to sun. This suggests that in addition to TRMT61B, additional factors are involved in modulating the association of the m^1^A 947 16S rRNA modification with mtDNA gene expression, possibly acting in a tissue-dependent manner.

### Sun exposure likely impacts the association of the m^1^A 947 16S modification with altered mtDNA gene expression in human skin

Environmental conditions likely impact mtDNA gene expression. Indeed, UV irradiation was previously shown to alter mitochondrial gene expression in keratinocytes (Kelly and Murphy 2018). With this in mind, we considered our dataset of RNA-seq experiments from skin, comprising samples collected from two body sites of the same individuals, one sun-exposed and the other not exposed to sun. The two types of skin samples showed stark differences in terms of mtDNA gene expression, with skin not exposed to sun displaying significantly higher mtDNA transcript levels than matched sun-exposed samples (Fig. 6A). Strikingly, whereas the level of m^1^A 947 16S rRNA modification associated with altered gene expression in skin not exposed to sun, such association was not observed in sun-exposed skin (Fig. 6B).

**Fig. 6:**
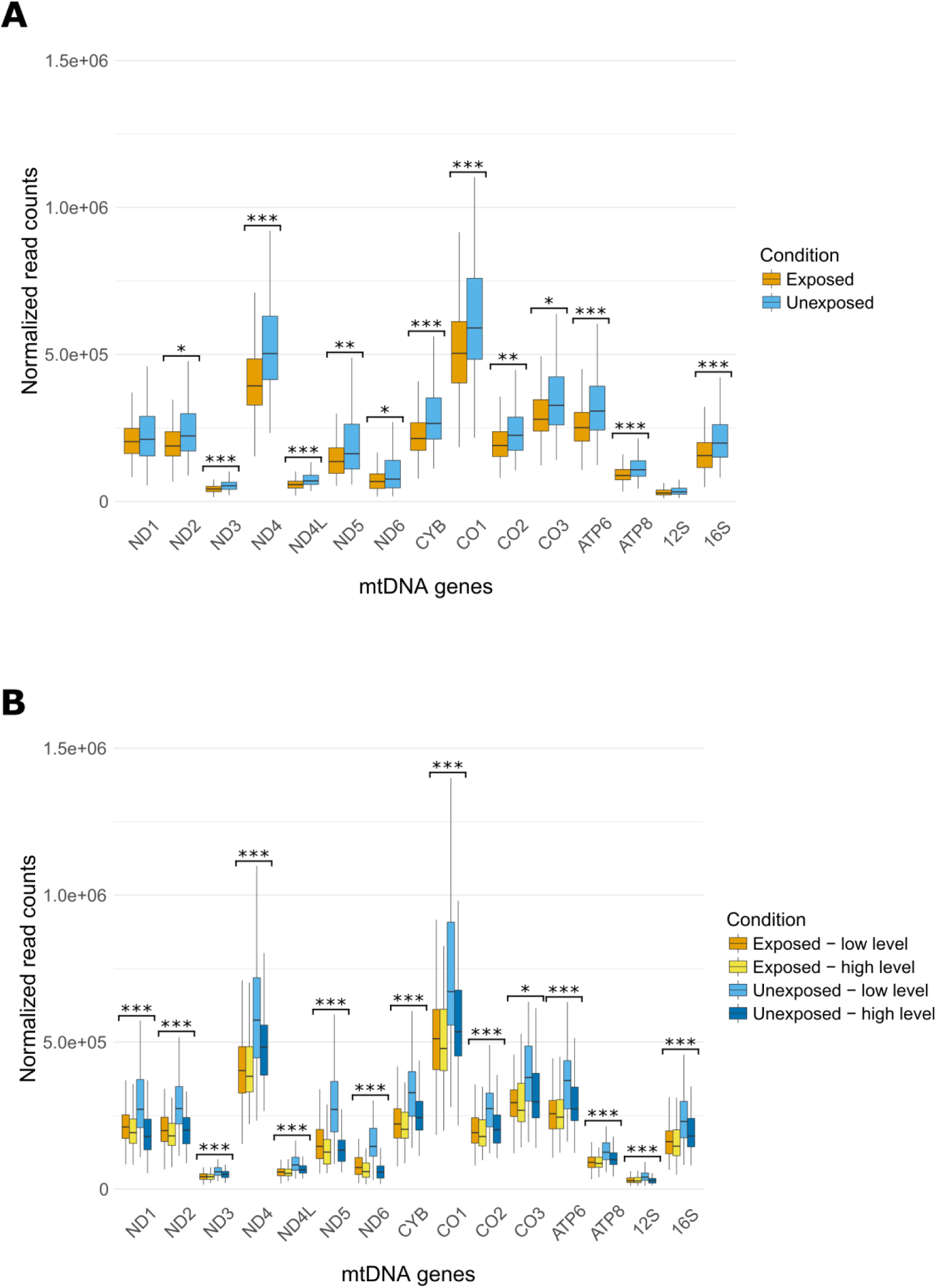
mtDNA gene expression differences between sun-exposed versus unexposed skin. (A) mtDNA gene expression levels were higher in skin not exposed to sun (blue), as compared to sun-exposed skin (orange). (B) m^1^A 947 16S rRNA modification levels associate with altered mtDNA gene expression in skin not exposed to sun but not in sun-exposed skin. Levels of the modification are indicated: In sun exposed skin - low (orange) and high (yellow), and in skin not exposed to sun - low (light blue) and high (blue). For both panel (A) and (B): X axis-mtDNA genes. Y axis-normalized read counts. Additionally, significance values are as follows: (*) < 3.3 × 10^−3^, (**) < 1 × 10^−4^, (***) < 1 × 10^−5^.

## Discussion

In the present study, we showed that both ancient and recurrent mtDNA SNPs associate with altered mtDNA transcript levels in a tissue-dependent manner, thus marking the identification of mtDNA eQTLs. Among those, recurrent mtDNA SNPs are the optimal candidates for causal eQTLs, as they are not consistently associated with a specific mtDNA genetic background, which could mask such functional potential. Secondly, the association of mt-eQTLs with tissue-dependent mtDNA gene expression raises the possibility that such expression patterns contribute to tissue-specific mitochondrial phenotypes (Boczonadi et al. 2018). This is consistent with previous reports of the involvement of ancient mtDNA SNPs in the penetrance of tissue-specific mitochondrial phenotypes, such as Leber hereditary optic neuropathy (LHON) (Brown et al. 1997; Torroni et al. 1997; Hudson et al. 2005). Indeed, close inspection of our analysis revealed that nearly half of the tissues, in which mt-eQTLs were discovered, are part of the nervous system and skeletal muscle, which are preferentially affected in mitochondrial disorders. Furthermore, the recurrent SNP at position 10398, which has been associated here with tissue-dependent mtDNA gene expression, had also been previously associated with altered tendency to develop a variety of phenotypes (Marom et al. 2017) such as breast cancer (Darvishi et al. 2007; Kulawiec et al. 2009) and type 2 diabetes (Rai et al. 2007). Therefore, since mtDNA transcription has not been systematically studied in the context of mitochondrial and complex disorders, our results offer a previously unexplored view on the underlying mechanism of such phenotypes.

The ancient and recurrent mt-eQTLs that were identified in the current study are located throughout the mtDNA sequence. Specifically, the recurrent mtDNA eQTLs, which are the best candidates to be causal, either occur immediately adjacent to a known regulatory element (position 263, near a preferential mitochondrial transcription factor A, TFAM, binding site), within the 12S rRNA gene (positions 750 and 1438) or within the ND3 gene (position 10398). This observation suggests that the impact of mtDNA SNPs on mtDNA transcription (Suissa et al. 2009) could be extended from the known non-coding regulatory elements to the entire mtDNA, and argues for the existence of novel regulatory elements throughout the mtDNA sequence. With this in mind, we recently identified *in-vivo* mtDNA binding of known regulators of nDNA gene expression within mtDNA protein-coding genes (Blumberg et al. 2014). Both findings imply that the mtDNA coding region is likely written not only in the protein-coding language, but also in the regulatory one. If this is true, one may consider a regulatory impact for certain SNPs occurring within genes. Once methods that enable manipulation of the mtDNA sequence become available, one will be able to test this hypothesis experimentally.

Our observed association between m^1^A 947 16S rRNA modification levels and differential mtDNA transcript levels in several tissues suggests the existence of regulatory interactions. Previously, we employed a bacterial model system to assess and demonstrate the importance of such modification for mitochondrial translation and cell life (Bar-Yaacov et al. 2016). Our current observation extends the role of the m^1^A 947 16S rRNA modification to association with mtDNA transcript levels in a variety of tissues. This suggests that mt-RNA modifications modulate gene expression but also, when specifically considering the m^1^A 947 16S rRNA modification, argues for a connection between mtDNA translation and transcript levels. The underlying mechanism of this connection should be further investigated in the future.

The observed bimodal distribution of m^1^A 947 16S rRNA modification levels in the human population across all tested tissues is intriguing. Apparently, one would have expected continuous distribution of the modified 16S rRNA level, with possible differences in the mean distribution among tissues. However, the complete absence of a 50±5% modification level across all tested body sites suggests the existence of an active modulation mechanism underlying the differentiation noted between higher and lower modification levels. If such a mechanism exists, then association of the modification levels with mtDNA gene expression may operate in an on-off manner.

Finally, our results suggest that the clear association of the m^1^A 947 16S rRNA modification with mtDNA transcript levels in skin not exposed to sun was lost in sun-exposed skin. Previously, it was shown that UV irradiation changes mitochondrial morphology and reduces mitochondrial oxygen consumption rate in skin keratinocytes, as well as altering expression of mitochondria-related genes (Leung et al. 2013; Schutz et al. 2016; Kelly and Murphy 2018). Accordingly, it is conceivable that the impact of m^1^A 947 16S rRNA modification on mtDNA transcript levels is influenced by UV exposure. This demonstrates a phenotypic outcome of such association, and lends support for the existence of a UV-sensitive mechanism that modulates the modification level.

In summary, we have shown that differential mtDNA gene expression associates with ancient and recurrent SNPs in different tissues, thus identifying for the first time, tissue-dependent mt-eQTLs. This is consistent with the previous suggestion that mitochondrial activity differs among tissues (Fernandez-Vizarra et al. 2011), and paves the way for future investigation of the association of such mt-eQTLs with tissue-specific mitochondrial phenotypes. Additionally, the occurrence of such eQTLs both in coding and non-coding mtDNA sequences imply the identification of previously overlooked regulatory elements. Secondly, since differential mtDNA expression associated with altered levels of the m^1^A 947 16S rRNA modification in eight tissues, mtDNA gene expression is likely modulated, at least in part, by this RNA modification. Such regulatory impact was particularly evident in sun-exposed skin, suggesting that this modulation of gene expression is sensitive to UV exposure. Taken together, the discovery of association of both mtDNA SNPs and an RNA modification with altered mtDNA gene expression, unearth a previously unexplored view on the impact of mtDNA variation on phenotypes, and lends new insight into the mechanisms underlying regulation of mtDNA gene expression.

## Methods

### GTEx RNA-seq and variant data

GTEx (Consortium 2013) v6 RNA-seq fastqs and whole genome VCFs were obtained from dbGaP (phs000424.v6.p1) (Mailman et al. 2007). Analysis was limited to tissues with data from at least 100 individuals (except for ovary), of whom at least 10% were black. Only the most recently collected samples were kept for replicate analysis.

### Mapping RNA-seq reads to human mtDNA

RNA-seq reads were trimmed using trimmomatic (Bolger et al. 2014) according to their fastQC and then mapped onto the entire human genome reference sequence (GRCh38) using STAR v2.5.3 (Dobin et al. 2013). The mapping process followed the ENCODE long mRNA protocol (https://github.com/ENCODE-DCC/long-rna-seq-pipeline), using the [--outFilterMismatchNoverLmax 0.05], [--alignSJoverhangMin 8], [--alignSJDBoverhangMin 1], [--alignSJDBoverhangMax 999] [--alignIntronMin 20], [--alignIntronMax 1000000], and [--alignMatesGapMax 1000000], with the modification of [--outFilterMultimapNmax 1] mapping parameters to achieve unique mapping. Since mtDNA sequence variability can impact the number of mapped RNA-seq reads, we re-mapped the reads against the same human genome files after replacing the mitochondrial reference sequence with the reconstructed mtDNA in an individual-specific manner. To this end, a revised index was generated for the new reference genomes by replacing the human mtDNA sequence with the reconstructed version, as previously described (Cohen et al. 2016). This was conducted separately for each of the tested samples, with the first-tier mapping files being retained. While generating the secondary mapping stage, most of the parameters of the first-tier mapping were retained while utilizing the 2-pass mode ([--twoPassMode Basic] and [--outFilterType BySJout]). Mapping accuracy was further increased by allowing fewer mismatches [--outFilterMismatchNmax 8], while analyzing couples of paired reads. To allow subsequent analysis of mtDNA gene expression in as many samples as possible we excluded samples with mtDNA read coverage of less than 165,690 (Supplemental Fig. S1). This approach enabled analyzing samples with mtDNA sequences covered by at least 10X across most mtDNA positions.

### Reconstruction of complete mtDNA sequences from RNA-seq data, multiple sequence alignment and phylogenetic analysis

Taking advantage of the polycistronic transcription of the mtDNA (Aloni and Attardi 1971), mtDNA sequences were reconstructed from each of the RNA-seq samples using an in-house script that followed the logic of the previously stablished MitoBamAnnotator (Zhidkov et al. 2011) tool. In brief, a pileup of the mtDNA-mapped reads was generated using the mpileup function in SAMtools (Li et al. 2009) against the revised mtDNA Cambridge reference sequence (rCRS) (Andrews et al. 1999). Bases with a phred score higher than 30 [-Q 30] were considered as a measure of quality control. The mapped bases were then counted, and the most frequent nucleotide in each mtDNA position was considered the major allele, only if that nucleotide position had a minimal coverage of 10X. To avoid strand bias, the secondary mutation at a given nucleotide position was recorded only if it was represented by at least two reads per strand. To calculate the level of m^1^A 947 16S rRNA modification, the value was calculated only when a minimum coverage of 500X at mtDNA position 2617 was available, followed by calculation of the fraction of reads harboring thymidine at this position. Each reconstructed mtDNA sequence underwent haplogroup assignment using HaploGrep (Kloss-Brandstatter et al. 2011).

### Expression pattern analysis considering mtDNA SNPs

mtDNA sequences of all individuals were aligned to identify polymorphic positions. For each polymorphic position, the samples were divided into groups according to their allele assignment. Using the linear model implemented in the Matrix eQTL R package (Shabalin 2012) as previously described (Cohen et al. 2016), eQTL mapping was calculated according to allele assignment, while considering age, gender and cause-of-death by the Hardy index as covariates. Bonferroni correction for multiple testing was employed. To reduce the false positive discovery rate, we focused on SNPs shared by at least 5 individuals (Levin et al. 2013). To identify possible associations of nDNA-encoded genes with differential expression patterns of mtDNA genes, the analysis focused on known SNPs (from the GTEx exome-based VCF files) in the dataset of genes with known mitochondrial RNA-binding activity(Wolf and Mootha 2014; Cohen et al. 2016), in addition to transcription factors and RNA-binding proteins that were recently identified in human mitochondria but are not included in MitoCarta (i.e., c-Jun, JunD, CEBPb, Mef2D, KAT8, NFATC1, THRA) (Ardail et al. 1993; Fernandez-Vizarra et al. 2008; She et al. 2011; Blumberg et al. 2014; Lambertini et al. 2015; Chatterjee et al. 2016). To enable detection of possible correlations with sufficient statistical power, KAVIAR (Glusman et al. 2011) was used to focus on SNPs with NCBI reference IDs.

### Assessing co-expression of potential regulators of mtDNA gene expression with mtDNA transcripts

Co-expression of either of 116 nDNA-encoded potential regulators of mtDNA gene expression (Supplemental Table S4, Supplemental Dataset S3), with mtDNA gene expression was performed while employing Pearson correlation. In brief, we performed differential expression analysis of the above-mentioned nDNA-encoded genes in the context of our identified mtDNA SNP association with mtDNA gene expression. Then, co-expression was sought between the 15 mtDNA-encoded mRNA and rRNA transcripts and the nDNA genes that passed the differential expression cutoff (p<4.3 × 10^−4^) after Bonferroni-correction for multiple testing. The same approach was applied to our identified association between the m^1^A 947 16S rRNA modification and mtDNA gene expression.

## Supplemental Data

Supplemental Data Supplemental Data includes 5 tables, 2 figures and 3 datasets.

Supplemental Table S1: mtDNA haplotype assignment per individual.

Supplemental Table S2: Mapping statistics of samples per tissue.

Supplemental Table S3: 116 candidate regulators of the mtDNA gene expression.

Supplemental Table S4: Summary of nDNA genes that significantly co-expressed with the mtDNA genes in the context of the identified recurrent mt-eQTLs.

Supplemental Table S5: Summary of nDNA genes that significantly co-expressed with the mtDNA genes in the context of the m^1^A 947 16S rRNA modification.

Supplemental Fig. S1: mtDNA positions covered by the RNA-seq reads across tissues.

Supplemental Fig. S2: mtDNA gene expression patterns differ between individuals with low vs. high m^1^A 947 16S rRNA modification levels.

Supplemental Dataset S1: eQTL analysis of mtDNA SNP association with mtDNA gene expression per tissue (including output of the MATRIX eQTL tool). Highlighted are the significant p-values after Bonferroni correction.

Supplemental Dataset S2: eQTL analysis of nDNA SNP association with mtDNA gene expression per tissue (including output of the MATRIX eQTL tool). Highlighted are the significant p-values after Bonferroni correction.

Supplemental Dataset S3: Results of co-expression analysis of 116 candidate regulators and mtDNA gene expression in the contexts of haplogroup-defining SNPs, recurrent SNPs and the m^1^A 947 16S rRNA modification.

## Acknowledgements

This study was funded by research grants from the Israel Science Foundation (ISF, 372/17), US-Israel Binational Science Foundation (2013060) and the US Army Life Sciences division (LS67993) awarded to DM. The authors thank the Negev Foundation for a scholarship for excellent graduate students awarded to TC.

## References

Aloni Y, Attardi G. 1971. Symmetrical in vivo transcription of mitochondrial DNA in HeLa cells. Proc Natl Acad Sci U S A 68: 1757–1761.

Andrews RM, Kubacka I, Chinnery PF, Lightowlers RN, Turnbull DM, Howell N. 1999. Reanalysis and revision of the Cambridge reference sequence for human mitochondrial DNA. Nat Genet 23: 147.

Ardail D, Lerme F, Puymirat J, Morel G. 1993. Evidence for the presence of alpha and beta-related T3 receptors in rat liver mitochondria. Eur J Cell Biol 62: 105–113.

Bar-Yaacov D, Avital G, Levin L, Richards AL, Hachen N, Rebolledo Jaramillo B, Nekrutenko A, Zarivach R, Mishmar D. 2013. RNA-DNA differences in human mitochondria restore ancestral form of 16S ribosomal RNA. Genome Res 23: 1789–1796.

Bar-Yaacov D, Frumkin I, Yashiro Y, Chujo T, Ishigami Y, Chemla Y, Blumberg A, Schlesinger O, Bieri P, Greber B et al. 2016. Mitochondrial 16S rRNA Is Methylated by tRNA Methyltransferase TRMT61B in All Vertebrates. PLoS Biol 14: e1002557.

Barshad G, Blumberg A, Cohen T, Mishmar D. 2018a. Human primitive brain displays negative mitochondrial-nuclear expression correlation of respiratory genes. Genome Res 28: 952–967.

Barshad G, Marom S, Cohen T, Mishmar D. 2018b. Mitochondrial DNA Transcription and Its Regulation: An Evolutionary Perspective. Trends in genetics : TIG 34: 682–692.

Blumberg A, Sailaja BS, Kundaje A, Levin L, Dadon S, Shmorak S, Shaulian E, Meshorer E, Mishmar D. 2014. Transcription factors bind negatively-selected sites within human mtDNA genes. Genome Biol Evol 6: 2634–2646.

Boczonadi V, Ricci G, Horvath R. 2018. Mitochondrial DNA transcription and translation: clinical syndromes. Essays Biochem doi:10.1042/EBC20170103.

Bolger AM, Lohse M, Usadel B. 2014. Trimmomatic: a flexible trimmer for Illumina sequence data. Bioinformatics 30: 2114–2120.

Brown MD, Sun F, Wallace DC. 1997. Clustering of Caucasian Leber hereditary optic neuropathy patients containing the 11778 or 14484 mutations on an mtDNA lineage. American journal of human genetics 60: 381–387.

Calvo SE, Clauser KR, Mootha VK. 2016. MitoCarta2.0: an updated inventory of mammalian mitochondrial proteins. Nucleic Acids Res 44: D1251–1257.

Chatterjee A, Seyfferth J, Lucci J, Gilsbach R, Preissl S, Bottinger L, Martensson CU, Panhale A, Stehle T, Kretz O et al. 2016. MOF Acetyl Transferase Regulates Transcription and Respiration in Mitochondria. Cell 167: 722–738 e723.

Cohen T, Levin L, Mishmar D. 2016. Ancient Out-of-Africa Mitochondrial DNA Variants Associate with Distinct Mitochondrial Gene Expression Patterns. PLoS Genet 12: e1006407.

Consortium GT. 2013. The Genotype-Tissue Expression (GTEx) project. Nat Genet 45: 580–585.

Darvishi K, Sharma S, Bhat AK, Rai E, Bamezai RN. 2007. Mitochondrial DNA G10398A polymorphism imparts maternal Haplogroup N a risk for breast and esophageal cancer. Cancer Lett 249: 249–255.

Dobin A, Davis CA, Schlesinger F, Drenkow J, Zaleski C, Jha S, Batut P, Chaisson M, Gingeras TR. 2013. STAR: ultrafast universal RNA-seq aligner. Bioinformatics 29: 15–21.

Fernandez-Vizarra E, Enriquez JA, Perez-Martos A, Montoya J, Fernandez-Silva P. 2008. Mitochondrial gene expression is regulated at multiple levels and differentially in the heart and liver by thyroid hormones. Current genetics 54: 13–22.

Fernandez-Vizarra E, Enriquez JA, Perez-Martos A, Montoya J, Fernandez-Silva P. 2011. Tissue-specific differences in mitochondrial activity and biogenesis. Mitochondrion 11: 207–213.

Glusman G, Caballero J, Mauldin DE, Hood L, Roach JC. 2011. Kaviar: an accessible system for testing SNV novelty. Bioinformatics 27: 3216–3217.

Gomez-Duran A, Pacheu-Grau D, Lopez-Gallardo E, Diez-Sanchez C, Montoya J, Lopez-Perez MJ, Ruiz-Pesini E. 2010. Unmasking the causes of multifactorial disorders: OXPHOS differences between mitochondrial haplogroups. Human molecular genetics 19: 3343–3353.

Hodgkinson A, Idaghdour Y, Gbeha E, Grenier JC, Hip-Ki E, Bruat V, Goulet JP, de Malliard T, Awadalla P. 2014. High-resolution genomic analysis of human mitochondrial RNA sequence variation. Science 344: 413–415.

Hudson G, Keers S, Yu Wai Man P, Griffiths P, Huoponen K, Savontaus ML, Nikoskelainen E, Zeviani M, Carrara F, Horvath R et al. 2005. Identification of an X-chromosomal locus and haplotype modulating the phenotype of a mitochondrial DNA disorder. American journal of human genetics 77: 1086–1091.

Kelly J, Murphy JE. 2018. Mitochondrial gene expression changes in cultured human skin cells following simulated sunlight irradiation. J Photochem Photobiol B 179: 167–174.

Kenney MC, Chwa M, Atilano SR, Falatoonzadeh P, Ramirez C, Malik D, Tarek M, Del Carpio JC, Nesburn AB, Boyer DS et al. 2014. Molecular and bioenergetic differences between cells with African versus European inherited mitochondrial DNA haplogroups: Implications for population susceptibility to diseases. Biochim Biophys Acta 1842: 208–219.

Kloss-Brandstatter A, Pacher D, Schonherr S, Weissensteiner H, Binna R, Specht G, Kronenberg F. 2011. HaploGrep: a fast and reliable algorithm for automatic classification of mitochondrial DNA haplogroups. Hum Mutat 32: 25–32.

Kulawiec M, Owens KM, Singh KK. 2009. mtDNA G10398A variant in African-American women with breast cancer provides resistance to apoptosis and promotes metastasis in mice. J Hum Genet.

Lambertini E, Penolazzi L, Morganti C, Lisignoli G, Zini N, Angelozzi M, Bonora M, Ferroni L, Pinton P, Zavan B et al. 2015. Osteogenic differentiation of human MSCs: Specific occupancy of the mitochondrial DNA by NFATc1 transcription factor. Int J Biochem Cell Biol 64: 212–219.

Leung MC, Rooney JP, Ryde IT, Bernal AJ, Bess AS, Crocker TL, Ji AQ, Meyer JN. 2013. Effects of early life exposure to ultraviolet C radiation on mitochondrial DNA content, transcription, ATP production, and oxygen consumption in developing Caenorhabditis elegans. BMC Pharmacol Toxicol 14: 9.

Levin L, Mishmar D. 2017. The genomic landscape of evolutionary convergence in mammals, birds and reptiles. Nature Ecology & Evolution 1: 0041.

Levin L, Zhidkov I, Gurman Y, Hawlena H, Mishmar D. 2013. Functional Recurrent Mutations in the Human Mitochondrial Phylogeny - Dual Roles in Evolution and Disease. Genome Biol Evol 5: 876–890.

Li H, Handsaker B, Wysoker A, Fennell T, Ruan J, Homer N, Marth G, Abecasis G, Durbin R, Genome Project Data Processing S. 2009. The Sequence Alignment/Map format and SAMtools. Bioinformatics 25: 2078–2079.

Mailman MD, Feolo M, Jin Y, Kimura M, Tryka K, Bagoutdinov R, Hao L, Kiang A, Paschall J, Phan L et al. 2007. The NCBI dbGaP database of genotypes and phenotypes. Nat Genet 39: 1181–1186.

Marom S, Friger M, Mishmar D. 2017. MtDNA meta-analysis reveals both phenotype specificity and allele heterogeneity: a model for differential association. Sci Rep 7: 43449.

Mishmar D, Ruiz-Pesini E, Brandon M, Wallace DC. 2004. Mitochondrial DNA-like sequences in the nucleus (NUMTs): insights into our African origins and the mechanism of foreign DNA integration. Hum Mutat 23: 125–133.

Rai E, Sharma S, Koul A, Bhat AK, Bhanwer AJ, Bamezai RN. 2007. Interaction between the UCP2-866G/A, mtDNA 10398G/A and PGC1alpha p.Thr394Thr and p.Gly482Ser polymorphisms in type 2 diabetes susceptibility in North Indian population. Human genetics 122: 535–540.

Schutz R, Kuratli K, Richard N, Stoll C, Schwager J. 2016. Mitochondrial and glycolytic activity of UV-irradiated human keratinocytes and its stimulation by a Saccharomyces cerevisiae autolysate. J Photochem Photobiol B 159: 142–148.

Shabalin AA. 2012. Matrix eQTL: ultra fast eQTL analysis via large matrix operations. Bioinformatics 28: 1353–1358.

She H, Yang Q, Shepherd K, Smith Y, Miller G, Testa C, Mao Z. 2011. Direct regulation of complex I by mitochondrial MEF2D is disrupted in a mouse model of Parkinson disease and in human patients. J Clin Invest 121: 930–940.

Sudmant PH, Alexis MS, Burge CB. 2015. Meta-analysis of RNA-seq expression data across species, tissues and studies. Genome Biol 16: 287.

Suissa S, Wang Z, Poole J, Wittkopp S, Feder J, Shutt TE, Wallace DC, Shadel GS, Mishmar D. 2009. Ancient mtDNA genetic variants modulate mtDNA transcription and replication. PLoS Genet 5: e1000474.

Torroni A, Petrozzi M, D’Urbano L, Sellitto D, Zeviani M, Carrara F, Carducci C, Leuzzi V, Carelli V, Barboni P et al. 1997. Haplotype and phylogenetic analyses suggest that one European-specific mtDNA background plays a role in the expression of Leber hereditary optic neuropathy by increasing the penetrance of the primary mutations 11778 and 14484. American journal of human genetics 60: 1107–1021.

Westra HJ, Franke L. 2014. From genome to function by studying eQTLs. Biochim Biophys Acta 1842: 1896–1902.

Wolf AR, Mootha VK. 2014. Functional Genomic Analysis of Human Mitochondrial RNA Processing. Cell reports 7: 918–931.

Zhidkov I, Nagar T, Mishmar D, Rubin E. 2011. MitoBamAnnotator: A web-based tool for detecting and annotating heteroplasmy in human mitochondrial DNA sequences. Mitochondrion 11: 924–928.

